# Which, when and where traits matter: functional trait-mediated species associations shift with microenvironmental and climatic variation

**DOI:** 10.64898/2026.01.21.700790

**Authors:** Ezequiel Antorán, Joaquín Calatayud, Ana L. Peralta, Adrián Escudero, Ana M. Sánchez, Arantzazu L. Luzuriaga, Marcelino de la Cruz

**Author notes:** Corresponding author: Ezequiel Antorán.

## Abstract

- Predicting coexistence in species-rich plant communities requires understanding the role of functional traits in species interactions and their resilience to environmental variation.
- In a manipulative factorial field experiment in an annual community (>45,000 individuals and 45 species), we followed plant associations from seedling to adult on both preserved and altered biological soil crust over two years with contrasting weather conditions. Neighborhood models quantified how among-species differences in key functional traits affected coexistence and how these effects varied in response to ontogenetic stage, environmental variation and spatial scale.
- Trait-based models revealed a prevalent negative relationship between functional differences and species associations, but these relationships varied predictively with stage, environment and spatial scale. Functional dissimilarity in traits such as specific leaf area, seed mass, and reproductive-to-vegetative biomass ratio consistently mediated species segregation, especially during wet years and among seedlings, while biological soil crust disturbance altered trait-mediated associations differently at fine and coarse spatial scales.
- Trait-mediated species associations shift predictably with environmental variation, suggesting that coexistence reflects a dynamic balance between environmental filtering and competitive interactions. This work provides novel experimental evidence of when, where, and which traits predict community assembly, which has implications for forecasting biodiversity responses to global environmental change.

## INTRODUCTION

Predicting species coexistence from functional traits remains one of the most challenging and unresolved questions in plant ecology. While functional traits have significantly advanced our understanding of ecosystem functioning and responses to environmental gradients, their predictive power regarding species interactions and coexistence remains limited (Westoby, 2025; Levine *et al*., 2025). This limitation arises in part because community dynamism and structure are profoundly shaped by a multitude of factors, including climatic conditions, environmental heterogeneity, and neighborhood interactions. Processes such as environmental filtering and competition leave imprints on the spatial organization of populations and communities (Seabloom *et al*., 2005; Biswas *et al*., 2016; Calatayud *et al*., 2020). Accordingly, spatial structure offers insights into the primary mechanisms driving the functioning and dynamics of ecological communities (McIntire & Fajardo, 2009; Borthagaray *et al*., 2014; Hart *et al*., 2017; Wiegand *et al*., 2021). One critical determinant of community spatial structure is the so-called *interspecific spatial association pattern*, where certain species pairs display non-random co-occurrence patterns (Wiegand *et al*., 2007; Wang *et al*., 2010; Keil *et al*., 2021), either positive or negative (i.e., they appear closer together or farther apart than expected under randomisation, respectively). Given that species’ patterns and responses to environmental conditions and neighborhood interactions are governed by their functional traits (Pescador *et al.,* 2018; López-Angulo *et al.,* 2021; López-Rubio *et al*., 2022; Ortiz *et al*., 2023) linking interspecific spatial association with the functional trait configuration of each species provides a tool for understanding the intricate processes underlying community dynamics (Kraft *et al*., 2015b; Ulrich *et al*., 2017; He & Biswas, 2019; Zambrano *et al*., 2020; Fajardo & Velázquez, 2021; Yin *et al*., 2021; Chalmandrier *et al*., 2022; Peralta *et al*., 2023). This trait-mediated approach offers a pathway to predict when and where species are likely to coexist, helping to reveal the mechanistic basis of community assembly.

When referring to species within the same trophic guild, the sign of the relationship between pairwise spatial associations and trait dissimilarities can inform on the main mechanisms shaping community assembly. Positive relationships, where the more functionally dissimilar a pair of species is, the more spatially aggregated they appear (Fig. **1B, F**), which may reflect competitive exclusion among species exploiting the same resources. However, it is important to note that spatial segregation among similar species may also arise from contrasting responses to fine-scale environmental heterogeneity. Conversely, when two species show divergent resource requirements (i.e., patent differences in trait values), their competition lessens, allowing them to locally coexist, as explained by MacArthur & Levins (1967) in the limiting similarity hypothesis and Mayfield & Levine (2010) approach to competitive exclusion (but see Holden & Cahill 2024). Negative relationships between spatial association and functional dissimilarity, i.e., the more similar the pair of species are, the more they aggregate together (Fig. **1**) suggest that specific trait values determine which species can spatially coexist within the community. This has been classically explained as environmental filtering (Bernard-Verdier *et al*., 2012; Cadotte & Tucker, 2017; Pescador *et al.,* 2018; López-Angulo *et al.,* 2021; Veresoglou *et al.,* 2024), where environmental or habitat processes act as a “filter” of species lacking appropriate traits to survive particular environmental conditions. Another explanation for the aggregation of functionally similar species is trait-based competitive hierarchies (Fig. **1C, D**), whereby species sharing particular values for some trait (i.e., a competition trait) have a competitive advantage over species with different values, which would be excluded as inferior competitors (Chesson, 2000; Mayfield & Levine, 2010; Kunstler *et al*., 2012; Kraft *et al*., 2014; Lasky *et al*., 2014; Lai *et al*., 2015; Cadotte & Tucker, 2017; Carmona *et al*., 2019; Yin *et al*., 2021; Mahaut *et al*., 2023). It is relatively easy to distinguish whether environmental filtering or trait-based competitive hierarchies are responsible for the observed negative relationship between spatial association and functional dissimilarity, by comparing how interspecific spatial association responds to absolute (Fig. **1E**) or real (Fig. **1C**) trait differences (HilleRisLambers *et al*., 2012; Kunstler *et al*., 2012; Carmona *et al*., 2016; Yin *et al*., 2021).

**Fig. 1.**
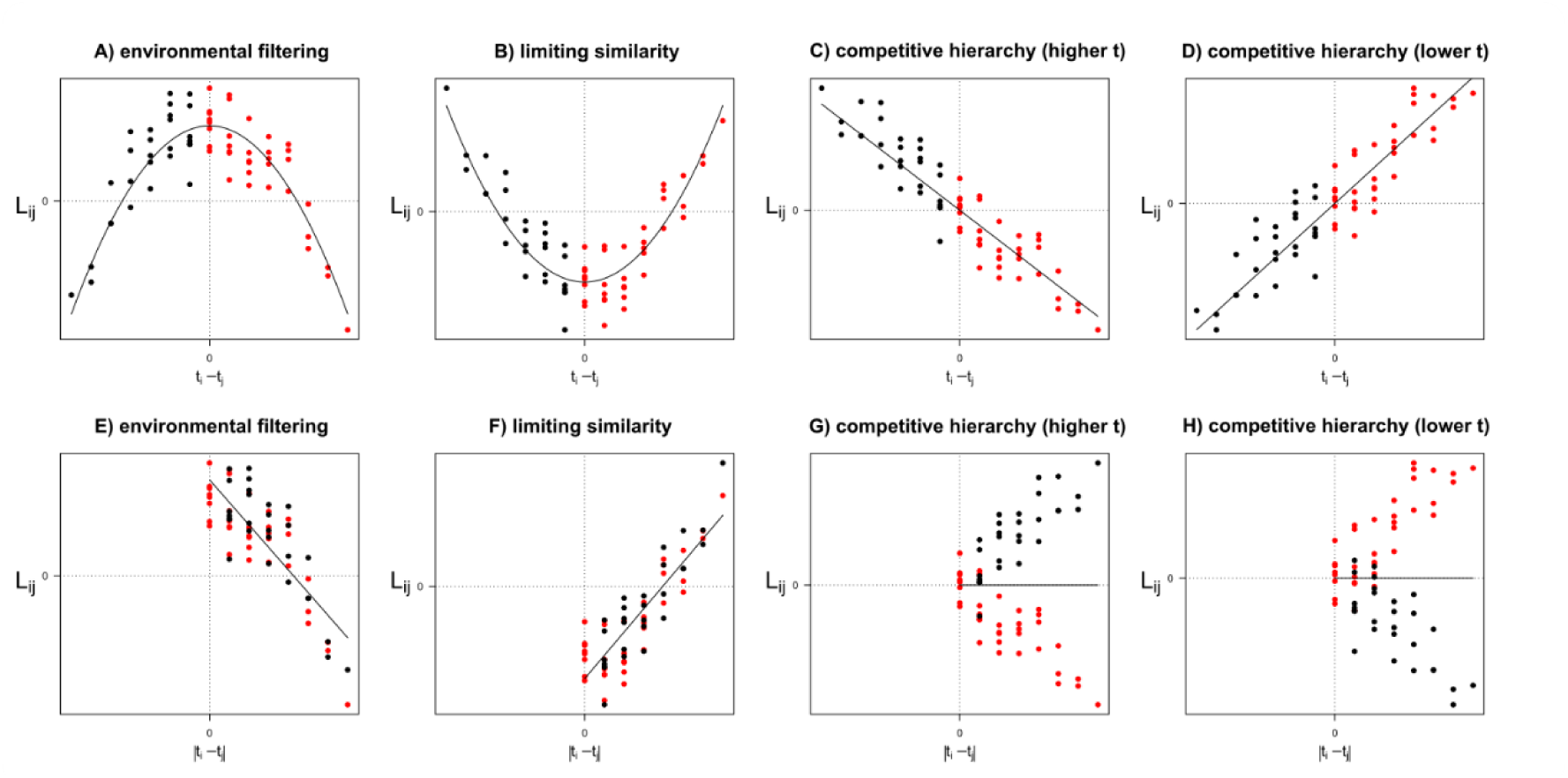
Conceptual framework illustrating the expected relationships between spatial association and trait dissimilarity between cooccurring species under different ecological processes. *L_ij_* is a normalized measure related to the average number of individuals of species *j* that coexist with each individual of species *i*. Values of *L_ij_* > 0 and *L_ij_* < 0 indicate respectively more and less neighbours than expected if both species are distributed independently from each other. *t_i_-t_j_* and | *t_i_-t_j_* | represent respectively the raw and absolute difference between the values of trait *t* for species *i* and *j*. Each point in the panels represents one pair of species. Under the process of “environmental filtering” it is expected that the more similar their values of *t* (i.e., the closer to 0 *t_i_-t_j_* and | *t_i_-t_j_* | are), the more spatially aggregated they will be (panels A, E). Contrarily, under “limiting similarity” it is expected that the more similar their values of *t*, the more spatially segregated (panels B, F). In contrast, under the hypothesis of “trait-based competitive hierarchies” species *j* will be more segregated from the vicinity of species *i* with increasing *t_i_-t_j_* values if the optimal is for large *t* values (panel C) or with decreasing *t_i_-t_j_* values if competitive ability is optimized for small *t* values (panel D). Panels g and h represent the expected null relationship between *L_ij_* and absolute trait differences and | *t_i_-t_j_* | under “trait-based competitive hierarchies”. Our statistical models do not directly test the mechanistic processes illustrated in panels A-D, but rather the expected patterns of spatial association consistent with those processes, as expressed by the relationships between *L* and trait differences. Black and red dots represent respectively negative and positive *t_i_-t_j_* values. The solid curves/lines represent linear models fitted to the data.

Despite the considerable attention given to this trait-based community assembly framework (HilleRisLambers *et al*., 2012; Kraft *et al*., 2015b; Carmona *et al*., 2016; Kunstler *et al*., 2016; Cadotte & Tucker, 2017; He & Biswas, 2019; Yin *et al*., 2021), the relationship between spatial associations and species’ trait dissimilarities is still not well understood (Davison *et al*. 2024, but see Bektaş *et al*. 2023). It remains unclear whether such relationships can vary over time, space, or environmental context (Westoby, 2025). For instance, changing climatic conditions can alter the importance of biotic interactions (*i.e*., competition *versus* facilitation) and environmental filtering in the assembly of plant communities (Maestre *et al*., 2009; He *et al*., 2013; Schleuning *et al*., 2020; Madrigal-González *et al*., 2023). Soil and light variation may also interfere with functional designs in complex ways (Li *et al*., 2021). Similarly, the importance of the difference between species may also shift throughout the ontogeny, showing opposing effects at different life stages (Gibert *et al*., 2016; Visser *et al*., 2016; Cooksley *et al*., 2024). Moreover, spatial associations may reflect not only species interactions but shared responses to subtle environmental gradients, underscoring the need for field approaches that combine manipulation with fine-scale spatial inference (Wiegand *et al*., 2007; Law *et al*., 2009; Cooksley *et al*., 2024). For example, in annual plant communities, a shift from early facilitation after germination at very fine spatial scales to strong competition in adult plants has been found (Schiffers & Tielbörger, 2006). In parallel, interactions with species from other trophic guilds can also modify assembly processes (Estes *et al*., 1998; Knight *et al*., 2005; Chesson & Kuang, 2008; Zhang *et al*., 2018). These interactions can exert direct effects, such as differential herbivory pressure modifying the species composition (Allbee *et al*., 2023), or indirect ones, such as the sorting of plant species due to the microenvironmental heterogeneity induced by biological soil crusts (BSC) (Belnap et al. 2016). In gypsum systems, BSCs strongly influence microsite conditions by altering soil water retention, infiltration, and nutrient availability (Chamizo *et al*., 2012; Escudero *et al*., 2015). These effects could determine germination success, seedling establishment, and subsequent competitive dynamics, especially for annual plants, which may be more responsive to fine-scale environmental variation due to their small size and rapid life cycle. Moreover, species could respond differently to BSC-driven heterogeneity, making BSCs a key factor to consider in community assembly. However, despite the well-known ecological influence of BSCs, their role in modulating trait-based assembly processes has rarely been tested experimentally. By explicitly manipulating BSC integrity, our approach allows us to disentangle whether BSC-driven microsite heterogeneity influences community assembly through environmental filtering or competitive interactions. Finally, assembly mechanisms can also vary with the effects of the spatial scale (Biswas *et al*., 2024), with pairwise species interactions exerting stronger effects at finer scales, while environmental conditions shape community responses at coarse scales (Hart *et al*., 2017). Overall, studying the variations in the intensity and sign of the relationship between spatial associations and traits differences between species under different conditions holds great potential to unravel assembly mechanisms, especially in the context of ongoing biodiversity redistribution and accelerating environmental change (Pecl *et al*., 2017).

Here, we used a field-based experimental design to assess how contrasting climatic conditions (rainfall), life stage, BSC condition, and spatial scale influence the assembly of a gypsum annual plant community. For this, we analyzed the relationships between interspecies spatial associations and species trait differences during two consecutive years with contrasting climatic conditions in terms of precipitation (wet *vs* dry), while experimentally manipulating the spatial heterogeneity of the BSC (intact vs disturbed BSC). This study encompasses two ontogenetic life stages (seedling *vs* adult) and spans different spatial scales. Within this framework, we aim to better understand the conditions under which functional traits can predict coexistence outcomes addressing two questions: 1) Does the sign and magnitude of the spatial association between pairs of species depend on their functional dissimilarities? And 2) Does the relationship between spatial associations and functional traits differences change in response to climatic conditions, BSC structure, ontogenetic stage, and spatial scale?

We expect that competition would force trait divergence for coexisting species at fine spatial scales (i.e., larger absolute differences in trait values for closer neighbor species). Conversely, at larger spatial scales, where environmental filtering processes may be prevalent, we expect spatial aggregation of functionally similar species. Finally, if a competition trait is involved, species associations will be related to the hierarchical distance of their trait values. In this context, if the spatial sorting of species is driven by competition, we expect a stronger positive relationship between spatial associations and functional trait dissimilarity in the wetter year where, under more benign conditions, competitive interactions should be stronger than in the dry, stressful year. This pattern is also anticipated to be more pronounced in adults and at smaller scales, where the signal of interactions should be more evident. Regarding BSC treatments, we anticipated that the intact BSC would provide greater environmental heterogeneity compared to the disturbed BSC, thereby generating microenvironmental filtering processes. Consequently, we expect a stronger negative relationship between spatial associations and trait dissimilarities in the intact plots. Altogether, our design enables a robust test of how competition, filtering, and trait differences interact to shape spatial organization under realistic field conditions.

## MATERIALS AND METHODS

### Study area

The experiment was carried out between summer 2013 and spring 2015, encompassing two consecutive annual plant life cycles in the El Espartal site (Valdemoro, Madrid, Central Spain, 40°11′11.5″N 3°37‘47.0″W – permit Ref. 10/058764.9/15 Comunidad de Madrid), specifically in a flat zone located in the upper section of a gypsum hill. Climate is characterized by a long-term average annual precipitation of 365 mm and a mean annual temperature of 15°C (www.aemet.es).

The vegetation in the study area is dominated by perennial shrubs, accounting for approximately 30% of the overall vegetation cover. Notably, many of these shrubs are specialized gypsum species, including *Helianthemum squamatum* (L.) Dum. Cours., *Lepidium subulatum* L., *Centaurea hyssopifolia* Vahl, and *Gypsophila struthium* L. in Loefl., as well as generalist species such as *Retama sphaerocarpa* (L.) Boiss., and *Macrochloa tenacissima* (L.) Kunth. These perennial species are distributed sporadically within a well-developed biological soil crust (BSC) matrix predominantly composed of lichens, namely *Diploschistes diacapsis* (Ach.) Lumbsch, *Squamarina lentigera* (G.H. Weber) Poelt, *Fulgensia subbracteata* (Nyl.) Poelt, and *Psora decipiens* (Hedw.) Hoffm (Luzuriaga *et al*., 2012).

The study site experiences seasonal coverage by a diverse annual plant community, with high plant densities reaching up to 4800 plants per square meter (Peralta et al., 2023). In years with abundant rainfall, this annual community can comprise up to 38 species within a 0.25 m^2^ area (Luzuriaga *et al*., 2012) and up to 100 species in only 45 m^2^ area (Luzuriaga *et al*., 2018). The annual plants in this community are characterized by their tiny size, with an average height of 9.7 ± 8.0 cm (Peralta et al. 2019), and most years they complete their life cycle between October and April. Among the most abundant annual species are *Asterolinon linum-stellatum* (L.) Duby in DC, *Bromus rubens* L., *Micropyrum tenellum* (L.) Link, *Helianthemum salicifolium* (L.) Mill., and *Ziziphora hispanica* L. Additionally, strict annual gypsophytes, including *Chaenorrhinum reyesii* (C. Vicioso & Pau) Benedí and *Campanula fastigiata* Dufour ex DC, are present in the study area.

### Experimental design and data collection

A total of six plots, each measuring 1 × 1 m, were randomly allocated within a flat area spanning approximately 0.4 ha. Care was taken to avoid small holes or depressions in the plot selection process, our primary focus was on the microenvironmental variations produced by the Biological Soil Crust (BSC). To ensure an adequate number of individuals per plot, one of the plots was extended to 1.20 × 1 m, aiming to accommodate a minimum of 1000 individuals. Among the six plots, three were designated as “intact BSC” plots, where the BSC layer remained undisturbed. Conversely, in the remaining three plots, the BSC was intentionally disrupted in the summer of 2013, before the first germination pulse, by carefully crushing the surface layer using a small mallet. The resulting fragments and powder were left on the soil surface, deliberately disrupting the physical continuity of the BSC without removing, homogenizing, or redistributing seeds, thereby creating what is referred to as the “disturbed BSC” condition (Peralta, 2018). In this way, the total soil seed bank was preserved, and the treatment specifically targeted the surface structure of the crust to test its role in spatial plant assembly. This manipulation reduced the fine-scale heterogeneity in surface roughness, infiltration, and nutrient micro-distribution generated by the intact crust, thus creating two contrasting conditions of microenvironmental variability.

Plant monitoring was conducted during two consecutive years, encompassing two complete life cycles of the annual plants. The first year, autumn precipitation was 33% below the average (herein, dry year), while the second year experienced a 35% above-average rainfall (i.e., wet year). Within each year, data collection occurred twice to capture the spatial patterns of plants in both the seedling (December) and adult (April) stages; totaling four sampling events per plot (two years × two plant life stages). We documented every emerging plant of each species, amounting to a total of 45,571 individuals and 45 species, across the six studied plots. We identified each plant and marked its rooting position on DIN-A3 transparent vinyl sheets made of polyvinyl chloride (PVC). To prevent any damage to the plants and biocrust, a mobile transparent methacrylate structure was employed, suspended 10 cm above the plot. After recording the data, the PVC sheets were converted into a single digital image per plot and spatial coordinates of the rooting point of each plant were captured using ArcGIS 10.1 software (ESRI, 2011). See Peralta, (2018) and Peralta et al. (2023) for details.

### Measuring functional trait dissimilarity

We measured functional traits for 25 of the 45 species recorded in our study area. For each species, we measured traits in a minimum of 10 healthy and undamaged adult individuals collected in the surroundings of our study area. Trait measurements were conducted in 2014 (dry year) following the established protocols described by Cornelissen et al. (2003) and Pérez-Harguindeguy et al. (2013) with further details provided in Peralta et al. (2019). We quantified specific leaf area (SLA), maximum plant height (H), specific root length (SRL), seed mass (SM), and the ratio of reproductive to vegetative biomass (RVR). SLA and H capture variation in aboveground resource use and light competition, whereas SRL captures belowground foraging capacity and adaptation to water availability. RVR integrates whole-plant biomass allocation and performance, and SM characterizes regeneration strategies under variable environmental conditions (Moles & Westoby, 2004; Hallett et al., 2011; Kunstler et al., 2012; Falster et al., 2018; Volaire et al., 2020). Together, these traits span the main axes of the plant economic spectrum (Díaz et al., 2016), and are mechanistically linked to competitive interactions and environmental filtering.

We computed two pairwise trait differences between species: absolute and hierarchical differences (Carmona *et al*., 2019; Yin *et al*., 2021). To quantify absolute differences (|*t_i_-t_j_*|) between species the average value pairs *i* and *j* for each trait *t*, we calculated the Euclidean distance *^t^D_ij_* based on each trait independently. Moreover, as an estimation of difference between species in the multivariate space of traits (i.e., all the considered traits together) we also computed a multitrait *^mt^D_ij_* as the absolute difference between scores of species *i* and *j* on the first component of a principal component analysis (PCA) of the matrix of standardized trait values. This first component represents a 41.2% of trait variance and summarizes the variation among SLA, SM, H and SRL, whereas the second axis (28.9% variance) was mainly related to the RVR and SLA traits (Fig. S1).

To quantify hierarchical differences (*t_i_-t^j^*), we computed hierarchical distance matrices *^t^H_ij_* as the difference of values for each trait *t* between species *i* and *j*. We also computed a multitrait *^mt^H_ij_* distance matrix computed as the difference between scores of species *i* and *j* on the first component of the above mentioned multitrait PCA (Yin *et al*., 2021).

### Measuring species interspecific spatial associations

We recorded a total of 45 annual plant species over the two study years. However, we focused only on the 25 species that had a minimum of 15 individuals per plot and sampling date. These species accounted for the vast majority of individuals in the community (97.22% across plots and sampling dates), providing sufficient replication for robust estimation of interspecific trait differences. We acknowledge that this approach underrepresents rare species, a common limitation in trait-based analyses (Carmona et al., 2016; Peralta et al., 2023), but the analyzed species capture the functional core of the community. Hereafter, we refer to the spatial pattern of individuals of a species in a particular plot and sampling date as “species pattern”.

### Test and meassurement of interspecific spatial associations

To characterize the interspecific spatial associations for each plot and sampling date, we employed the inhomogeneous cross-type *K_ij_* function, defined as:

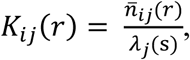

where, for two species *i* and *j*, *n̂ij (r)* is the expected number of neighbours of type *j* within a radius *r* around any point of type *i*, and *λj* (s) is the inhomogeneous intensity of the pattern of type *j* points (Baddeley et al. 2015). The intensity surfaces *λj* (s) were estimated by kernel smoothing (Gabriel *et al*., 2023). See details in the Supporting Information.

We computed *Kij* (r) for all pairs of species (among the 25 selected) in each plot and sampling date, and tested it against a null model of inhomogeneous Poisson independence (He & Biswas, 2019). This null model assumes that conditional on their inhomogeneous surfaces *λ_i_*(*s*) and *λ_j_*(*s*), the two patterns are independent. The test consists in comparing the *Kÿ* (r) function of the observed pattern with the envelopes obtained by simulating patterns from the null model, i.e., fixing the coordinates of pattern *i* and simulating *j* from *λ_j_*(*s*). The envelopes for each test were based on 39 simulations.

After computing the tests, we calculated for each *r* value the percentage of tests that showed significant aggregation, segregation or independence, i.e., the percentage of tests where the observed *K_ij_* (r) was respectively above, below or within the simulation envelopes (Fig. 2). This allowed us to identify two focal spatial scales: r = 40 mm, where significant positive associations were most frequent, and r = 250 mm, where significant negative associations predominated. These distances were selected using an objective, data-driven criterion based on the frequency of significant associations across distances. The 40 mm scale reflects the fine spatial range at which direct plant–plant interactions are most likely for small annual plants, whereas 250 mm represents a broader neighbourhood that integrates both interactions beyond immediate neighbours and microhabitat-related heterogeneity This larger scale is consistent with neighbourhood distances used in comparable studies of annual plant communities (e.g., Godoy & Levine, 2014; Bimler et al., 2018, 2024; Buche et al., 2025). Although 250 mm was the maximum feasible radius within our 1 m^2^ plots (Baddeley et al., 2015) and the proportion of negative associations continued to increase up to this limit (Fig. 2), it should be interpreted as the broadest scale accessible within our design rather than an upper bound for neighbourhood effects. For each plot and sampling date, we computed two spatial association matrices (one for each selected *r*) as *L_ij_*(r), i.e., the normalized version of *K_ij_* (r) (Besag 1977). These matrices measure the strength of spatial association between each pair of species *i* and *j*: if *L_ij_*(r) = 0 the two species are independent (at spatial scale *r*), if *L_ij_*(r) > 0 or *L_ij_*(r) < 0 the two species are positively or negatively associated, respectively, and the larger is the absolute value *L_ij_* (r), the stronger the association.

**Fig. 2.**
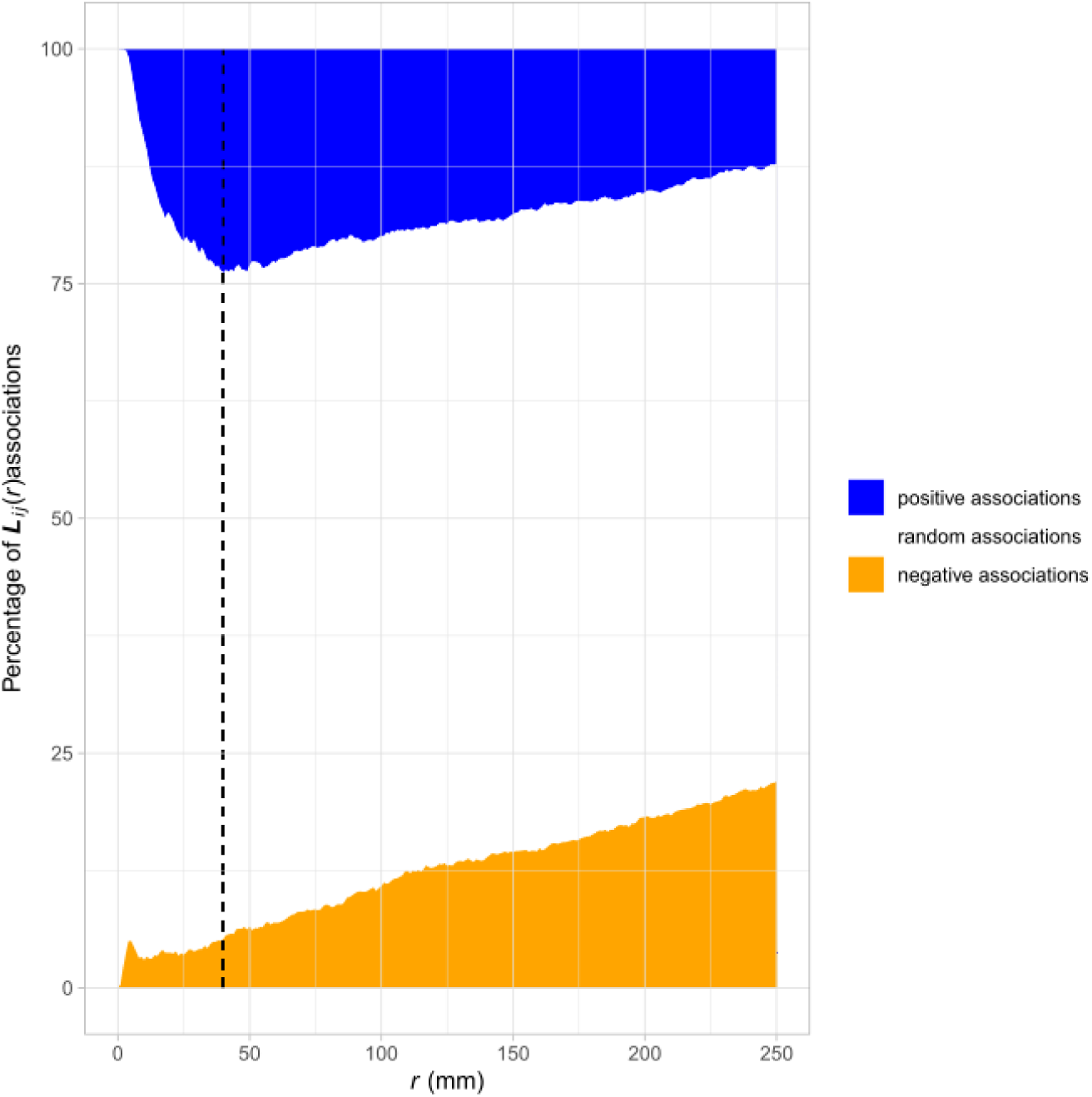
Variation in the percentage of significant (positive and negative) tests of interspecific spatial association (*L_ij_* (*r*)) against null models of inhomogeneous Poisson independence with the spatial scale *r*. The black, dashed vertical line indicates the spatial scale (*r* = 40 mm) where the largest proportion of significant positive associations was found. The largest proportion of significant negative associations was found for the largest spatial scale considered, i.e., *r* = 250 mm.

In total, we obtained 48 *L_ij_* (r) matrices of inter-species spatial association, i.e., six plots × two years × two plant life stages × two spatial scales. It is important to note that the number of species pairs (i.e., matrix dimension) varied in each plot and sampling date, as we only considered species with a minimum of 15 individuals but, in total, they amounted to 2492 species pairs.

### Relationship between species functional dissimilarity and species spatial association

To compare the relative effect of absolute *vs* hierarchical trait distances on the spatial association of species across different treatments, we fitted linear mixed models (one for each trait *t* and also for the multitrait distances) which included as predictors both absolute (*^t^D_ij_*) and hierarchical (*^t^H_ij_*) distances and their interactions with the experimental factors (BSC, year and life-stage). Because the same species appear in multiple species pairs, we included random intercept and slope effects of species *j* (*sp_j_*) nested within species *i* (*sp_i_*); we included also a random intercept for plot (Carmona *et al*., 2019; Yin *et al*., 2021). The models take the general form:

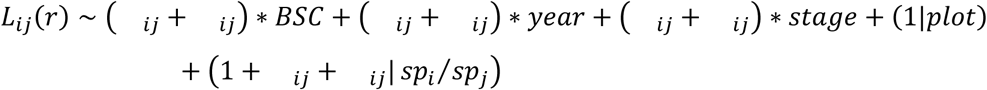

Note that although the experiment consisted of six 1 m^2^ plots (three per BSC treatment), our unit of inference was not the plot itself but the pairwise spatial associations between species. Each plot contained tens of species and thousands of individual plants, generating hundreds of species pair observations per plot (also per treatment, life stage, and year. Across all combinations of BSC treatment, life stage, and year (n = 24 plot-level contexts), we obtained >2,400 species-pair replicates involving >44,300 individuals from 25 species.

All analyses were conducted in the R software (R Core Team 2025). The PCA to summarize trait values was computed with function rda() of “vegan” package (Oksanen et al. 2022). Spatial associations were computed with function Kcross.inhom() in package “spatstat” (Baddeley *et al*., 2015). Linear mixed models were fitted and assessed using functions glmmTMB() of the “glmmTMB” package (Brooks *et al*., 2017) and Anova() of package “car” (Fox & Weisberg, 2019).

## RESULTS

Our linear mixed models showed that most effects on interspecific spatial associations were mediated by absolute (*^t^D_ij_*) rather than hierarchical (*^t^H_ij_*) functional distances (Tables 1 and 2), with a prevalent negative relationship, i.e. higher trait dissimilarity was related to lower spatial aggregation of species (Fig. 3 and Fig. 4). At fine scale (*r* = 40 mm), the effects of functional distances depended either on the life stage or climate for most functional traits (Table 1). For SLA, H and multitrait combination, the negative relationship between *L_ij_* (r) and *^t^D_j_* was stronger during the second (i.e., wet) year (*p* < 0.05; Table 1, Table S1). During the wet years, smaller functional differences favoured closer aggregations whereas larger functional differences favoured more segregation than during the dry year (Fig. 3). In the case of SLA, RVR and multitrait dissimilarities, the relationship was also stronger for seedlings than for adults (Table 1, Fig. 3), with larger functional differences favouring stronger spatial segregation in seedlings than in adults. The BSC treatment showed discernible effects only for SM, with a stronger negative relationship between *L_ij_* (*r*) and *^t^D_j_* in the disturbed than in the intact BSC treatment (*p* < 0.05).

**Fig. 3.**
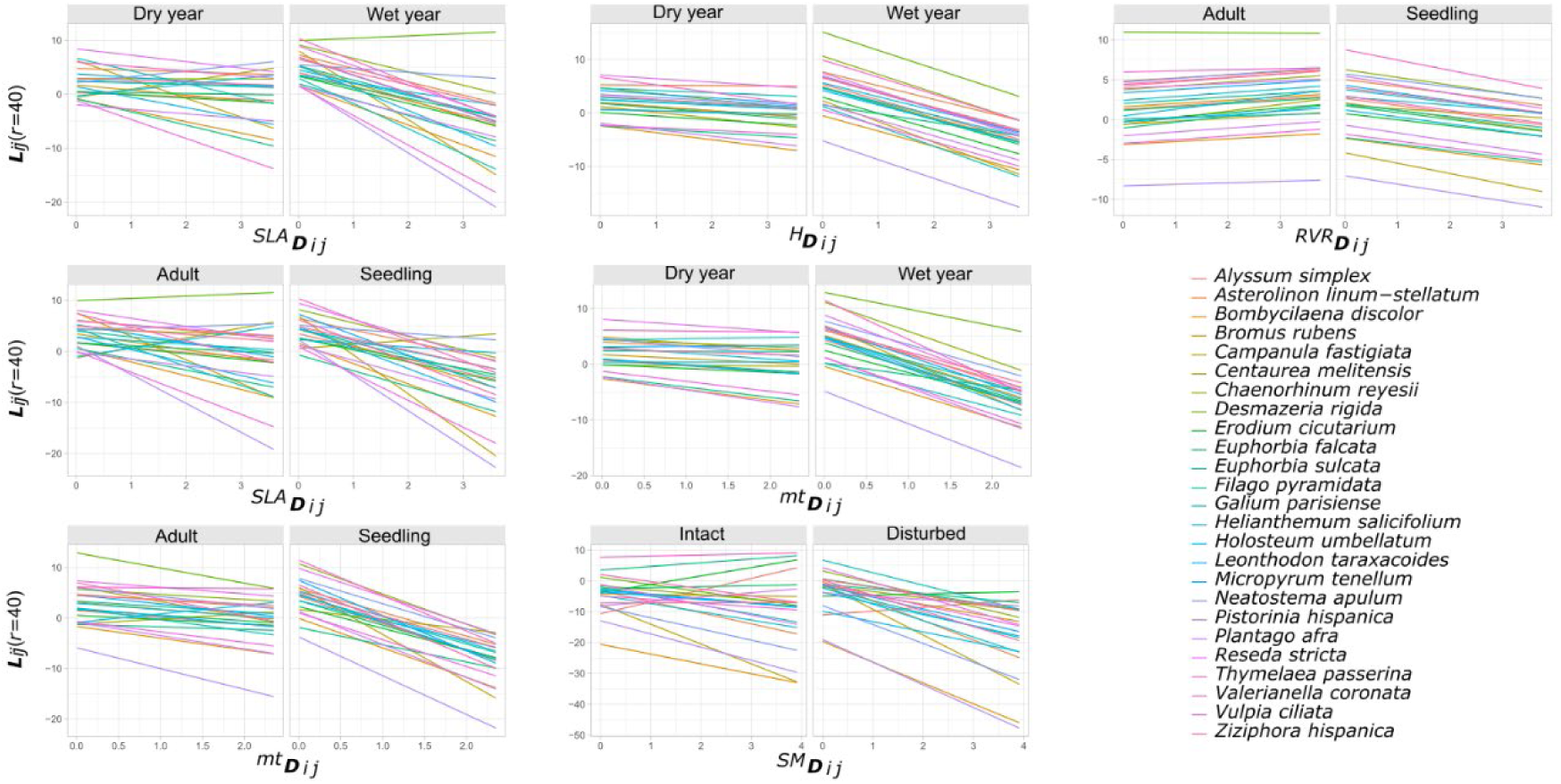
Fitted relationship between spatial association *L_ij_*at the coarse spatial scale (*r*=40 mm) and trait dissimilarity (*^t^D_ij_*) for some of the significant effects summarized in Table 1. The acronyms for *t* in *^t^D_ij_* are SLA: specific leaf area; H: maximum plant height; RVR: reproductive-vegetative ratio; mt: combination of all traits (“multitrait”). “Dry year” and “Wet year” correspond to the two selected years of sampling. “Intact” and “Disturbed” are the two treatments for the biological soil crust (BSC). “Adult” and “Seedling” correspond to the life stages.

**Fig. 4.**
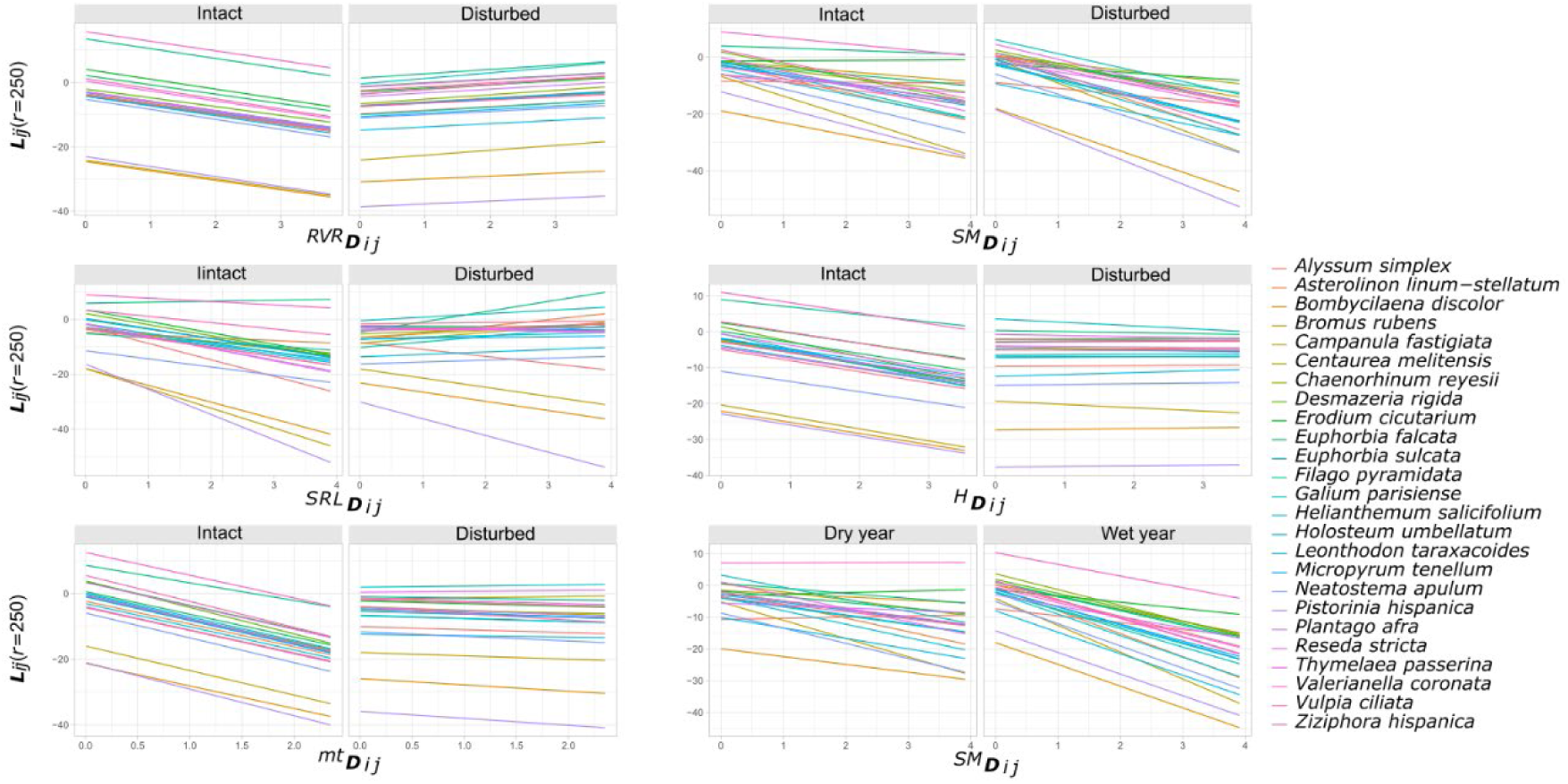
Fitted relationship between spatial association *L_ij_*at the coarse spatial scale (*r=*250 mm) and absolute trait disimilarity (*^t^D_ij_*) for some of the significant effects summarized in Table 2. The acronyms for *t* in *^t^D_ij_* are SLA: specific leaf area; SM: seed mass; H: maximumplant height; RVR: reproductive-vegetative ratio; mt: combination of all traits (“multitrait”). “Dry year” and “Wet year” correspond to the two selected years of sampling. “Intact” and “Disturbed” are the two treatments for the biological soil crust (BSC). “Adult” and “Seedling” correspond to the life stages.

**Table 1.** Summary of the results of linear mixed models (including “plot” and species *j* nested within species *i* as random effects) explaining interspecific spatial association *L_ij_* at the fine spatial scale (*r*= 40 mm) as a function of absolute (*^t^D_ij_*) and hierarchical (*^t^H_ij_*) trait distances and years, life-stage (Stage) and biological soil crust (BSC) treatments for the functional traits (*t*) evaluated (SLA: specific leaf area; RVR: reproductive-vegetative ratio; SM: seed mass; SRL: Specific root length; H: maximum plant height; mt: multitrait, combination of all traits). Values are Chi-square statistics (df = 1) with significance codes as superscripts: *** p < 0.001; ** p < 0.01; * p < 0.05; p < 0.1; – p > 0.1. Detailed results are provided in Table S1.

| Term | Trait $t$ | | | | | |
| --- | --- | --- | --- | --- | --- | --- |
|  | SLA | RVR | SM | SRL | H | mt |
| ${}^tD_{ij}$ | 9.30 ** | 0.07 – | 2.55 – | 0.00 – | 14.92 *** | 9.62 ** |
| ${}^tH_{ij}$ | 0.01 – | 1.18 – | 0.00 – | 0.37 – | 0.58 – | 0.15 – |
| Stage | 0.86 – | 1.06 – | 3.15 • | 1.09 – | 1.31 – | 1.15 – |
| BSC | 0.13 – | 0.18 – | 0.00 – | 0.19 – | 0.17 – | 0.13 – |
| Year | 5.41 * | 3.58 • | 1.37 – | 4.51 * | 3.90 * | 3.58 • |
| ${}^tD_{ij} \times \text{stage}$ | 5.13 * | 4.93 * | 1.14 – | 0.02 – | 0.01 – | 14.88 *** |
| ${}^tH_{ij} \times \text{stage}$ | 0.01 – | 0.04 – | 0.00 – | 0.05 – | 0.00 – | 0.00 – |
| ${}^tD_{ij} \times \text{BSC}$ | 0.62 – | 1.43 – | 7.36 ** | 0.10 – | 2.92 • | 0.79 – |
| ${}^tH_{ij} \times \text{BSC}$ | 0.02 – | 0.10 – | 2.29 – | 0.16 – | 0.00 – | 0.00 – |
| ${}^tD_{ij} \times \text{year}$ | 9.13 ** | 0.09 – | 3.15 • | 0.01 – | 10.57 ** | 11.70 *** |
| ${}^tH_{ij} \times \text{year}$ | 0.03 – | 0.07 – | 0.38 – | 0.00 – | 0.09 – | 0.01 – |

**Table 2.** Summary of the results of linear mixed models (including “plot” and species *j* nested within species *i* as random effects) for explaining interspecific spatial association *L_ij_* at the coarse spatial scale (*r*= 250 mm) as a function of absolute (*^t^D_ij_*) and hierarchical (*^t^**H**_ij_*) trait distances and years, life-stage (Stage) and biological soil crust (BSC) treatments for the functional traits (*t*) evaluated (SLA: specific leaf area; RVR: reproductive-vegetative ratio; SM: seed mass; SRL: Specific root length; H: maximum plant height; mt: multitrait, combination of all traits). Values are Chi-square statistics (df = 1) with significance codes as superscripts: *** p < 0.001; ** p < 0.01; * p < 0.05; • p < 0.1; – p > 0.1. Detailed results are provided in Table S2.

| Term | Trait $t$ | | | | | |
| --- | --- | --- | --- | --- | --- | --- |
|  | SLA | RVR | SM | SRL | H | mt |
| ${}^tD_{ij}$ | 7.65 ** | 0.44 – | 11.98 *** | 1.63 – | 2.58 – | 4.10 * |
| ${}^tH_{ij}$ | 0.00 – | 9.52 ** | 0.53 – | 1.41 – | 0.22 – | 0.23 – |
| Stage | 2.76 • | 2.90 • | 3.15 • | 2.65 – | 2.57 – | 2.78 • |
| BSC | 0.00 – | 0.00 – | 0.00 – | 0.01 – | 0.01 – | 0.00 – |
| Year | 0.43 – | 0.55 – | 0.31 – | 0.33 – | 0.57 – | 0.40 – |
| ${}^tD_{ij} \times \text{stage}$ | 1.70 – | 0.57 – | 1.17 – | 0.16 – | 3.79 • | 0.03 – |
| ${}^tH_{ij} \times \text{stage}$ | 0.05 – | 0.00 – | 0.02 – | 0.09 – | 0.09 – | 0.06 – |
| ${}^tD_{ij} \times \text{BSC}$ | 0.45 – | 14.34 *** | 4.79 * | 8.65 ** | 8.06 ** | 10.20 ** |
| ${}^tH_{ij} \times \text{BSC}$ | 0.87 – | 0.03 – | 1.32 – | 0.05 – | 0.01 – | 0.33 – |
| ${}^tD_{ij} \times \text{year}$ | 0.03 – | 0.17 – | 5.73 * | 1.21 – | 0.01 – | 1.63 – |
| ${}^tH_{ij} \times \text{year}$ | 0.43 – | 0.00 – | 0.17 – | 0.04 – | 0.74 – | 0.43 – |

At coarse scale (*r* = 250 mm), the effects of absolute differences (*^t^D_ij_*) depended mostly on the BSC treatment (Table 2). For example, the effects of RVR, SRL, H and multitrait functional distances were inexistent or less pronounced in the disturbed treatment, whereas the contrary occurred for SM (i.e., more negative effects of *^SM^D_ij_* in the disturbed treatment) (Fig. 4). The negative effect of *^SLA^D_j_* on *L_ij_* did not vary among BSC treatments (Table 2). In the case of SM, the relationship was also stronger during the wet year (*p* < 0.05; Table 2). At this coarse scale we found the only significant effect of hierarchical differences (*^RVR^H_ij_*) on the spatial association between species (Table 2 and Fig. S2).

## DISCUSSION

Most studies assessing the role of functional traits in community assembly have analyzed the convergence or divergence of trait values at plot scale throughout long environmental gradients (e.g., Swenson & Enquist, 2007; Cornwell & Ackerly, 2009; Spasojevic & Suding, 2012; Lhotsky *et al.,* 2016; López-Angulo *et al.,* 2021; Bektaş *et al*., 2023). Recent studies show that trait dissimilarity between plants can also explain the sign and magnitude of interspecies spatial associations within communities (de Bello et al. 2013; He & Biswas, 2019). Our study goes a step further and shows that the effect of trait differences on species associations and assembly is not fixed, but varies depending on variations in BSC integrity, climatic conditions and plant life stage. This suggests that assembly mechanisms related to functional traits may be more labile than previously thought. Moreover, this flexibility indicates that increasing the predictive power of functional traits models (which) for explaining species coexistence requires explicit consideration of spatial scale and abiotic heterogeneity (i.e., where) and both ontogenetic stage and climatic conditions (i.e., when). Our interpretations rely on the well-established assumption that emergent spatial patterns can provide indirect evidence of underlying biotic and abiotic processes (McIntire & Fajardo, 2009; Wiegand & Moloney, 2013). Therefore, while spatial associations alone do not constitute direct proof of mechanism, the consistency of trait–association relationships across years, life stages, and BSC treatments supports the robustness of our inferences.

We found some support for the hypothesis that pairwise competition between species is linked to hierarchical distances in the ratio of reproductive to vegetative biomass (RVR) This was the only trait where a hierarchical effect (Kunstler et al. 2012), was detected (Fig. S2). This trait integrates biomass allocation and plant performance across ontogeny, and traits related to whole-plant allocation have been shown to capture responses to environmental variation more strongly than other functional traits (Falster *et al*., 2018; Volaire *et al*., 2020) and it may be considered a summary of the competitive ability of plants. Although both the large scale involved (i.e., *r* = 250 mm) and the negative relationship between spatial association and RVR differences could suggest the involvement of environmental filtering, the fact that it is the raw and not the absolute difference in RVR values which is negatively related to spatial association suggest a trait-based competitive hierarchy. This negative effect of interspecific RVR differences on spatial associations was irrespective of year, life stage or BSC treatment. This means that competitive ability is associated to higher RVR values (Fig. **1C**). In fact, the higher values for this trait are for opportunistic species (e. g., *Leontodon taraxacoides* and *Desmazeria rigida*), which can cope with a wide range of resource availability conditions and are, therefore, insensible to BSC structure (Peralta et al. 2023). This allows them to germinate and establish without restrictions of any kind in the microsites where their seeds were primarily dispersed, usually around the mother plant, thereby generating clustered patterns which may hamper the establishment of dissimilar species. This high biomass allocation to reproduction (high RVR) likely reflects a strategy of high fecundity and reproductive economy (Aarssen et al., 2006; Tracey & Aarssen, 2011 allowing opportunistic species to saturate local microsites. In fact, Peralta et al. (2023) found that opportunistic species with high RVR values exhibited clustered homogeneous spatial patterns (i.e. not affected by environmental heterogeneity), whereas conservative species (low RVR) mainly had non-clustered heterogeneous spatial patterns, suggesting a greater dependence on the microenvironmental heterogeneity provided by the BSC rather than on offspring number. Note, however, that the absolute differences in RVR *(^RVR^DIĴ*) were also negatively related to the magnitude of interspecies negative associations, although only in the plots with intact BSC (Fig. 4), which suggest that, in addition to the trait-based competitive hierarchies, this trait is involved in the response to some kind of environmental filtering generated by the BSC.

Our results also highlight the primary importance of niche differences in structuring annual plant communities across all scales. The consistent negative relationship between spatial associations and absolute functional distances for the rest of traits considered, indicates a system where distinct microenvironmental preferences shape community composition. Such patterns suggest that coexistence in these communities is largely driven by environmental filtering rather than strong, asymmetric competition for resources. Under these conditions, where species performance depends more on microsite suitability than on size-dependent dominance, interactions tend to be more symmetric. This contrasts with the prevalent view that symmetric plant interactions are expected to arise only when competition comes from below-ground resources (Schwinning & Weiner, 1998; Mahaut *et al*., 2023). For example, at the larger scales (*r* = 250 mm), absolute functional distances for belowground traits, such as SRL, for aboveground traits such as H, and for all of them in the multitrait combination were negatively related to spatial associations (i.e., environmental filtering, Fig. **1E**) in the intact BSC plots, whereas in the disturbed plots they had no effect. This confirms the filtering effect exerted by BSC which affects the role of both belowground (SRL), aboveground (H) and whole plant (RVR, multitrait) traits in the assembly of this community. These results also agree with the well-known effects of the BSC in gypsum ecosystems, where it is a pivotal driver of microenvironmental heterogeneity (Escudero *et al*., 2015; Ortiz *et al*., 2023) by modulating water (Chamizo *et al*., 2012; Berdugo *et al*., 2014) and nutrient availability (Escudero *et al*., 2015; Belnap *et al*., 2016; Sánchez *et al*., 2022).

In the case of SLA, the negative relationship between the absolute functional distances and pairwise associations were maintained irrespective of BSC treatment, year or life stage (Table 2, Fig. 4). As SLA is related to resource use, this negative relationship suggests that other sources of environmental variation, in addition to BSC, are sorting species composition (at the same scales). Potential factors could include soil heterogeneity related to soil pH or organic content, as seen in Mediterranean dwarf-shrubland assemblages (Pescador *et al*., 2020) which could be mediated by differences in soil microbial activity at such scales (Han *et al*., 2023). Microbial influences on seed dormancy and germination, although not directly measured here, represent an underexplored mechanism and may be also driving these fine-scale assembly patterns.

We also found that the negative relationship between SM distances and spatial associations increased in BSC disturbed plots. SM is related to both establishment and survival of seedlings (Hallett *et al*., 2011) and to dispersal and acquisition of safe sites (Gonzalez & Ghermandi, 2012). It is plausible that once the main source of heterogeneity has been eliminated in the disturbed plots, other edaphic sources of heterogeneity like soil texture may filter which species enter the soil bank (and later germinate and establish as seedlings) on the basis of seed size (Benvenuti, 2007).

Negative relationships between spatial association and absolute trait differences have been commonly interpreted as environmental filtering (He & Biswas, 2019; Yin et al. 2021). We found, however, that these negative relationships were also patent at the small spatial scales (*r* = 40 mm) where biotic interactions, instead of environmental filtering, are expected to play a key role (Chesson, 2000; McGill, 2010). In addition, we found that for some traits (SLA, H, MT, and even SM) the negative relationship between absolute functional distances and spatial associations became stronger (i.e., increased the signal of environmental filtering) during the wet year. The fact that absolute ( *^t^D_ij_*) rather than hierarchical distances ( *^t^H_ij_*) were involved in these negative relationships suggest that the effect of micro-environmental heterogeneity is enhanced by favourable, wetter conditions. In fact, some authors have recognized that even at small spatial scales some within-site abiotic heterogeneity may contribute to a kind of environmental filtering (Kraft et al. 2015; Pescador et al. 2020). For example, some lichens (e.g., *Collema* spp.) and cyanobacteria in the BSC of our community are able to fix atmospheric N_2_ (Castillo-Monroy *et al*., 2010), which may increase soil fertility in their surroundings specially during wetter years. Lichens in the BSC differ also in their water holding capacity (Ghiloufi *et al*., 2023), which may temporarily affect processes where SLA, H or other functional traits (e.g., correlated with the multitrait) are involved, generating during the wetter years a stronger microheterogeneity within the general BSC heterogeneity in each plot. Since the effects of interactions and environmental preferences cannot be understood in isolation (Carvajal *et al*., 2023) and because assembly at small spatial scales is probably driven by a combination of both abiotic and biotic effects (e.g., Maire et al. 2012), our findings could be a consequence of shifting the average fitness of each species (Kraft et al. 2015) during the wet year. This could also explain the strong signals found in the wet year at small spatial scales, suggesting that biotic interactions would amplify the effects of environmental filtering.

Another interesting result was that SLA, RVR and especially the multitrait absolute distances were more negatively related to spatial associations at the seedling stage, i.e., that seedlings experienced a stronger environmental filtering for these traits. As expected, this occurred only at the fine spatial scale (*r* = 40 mm). As we commented before, those traits are related to resource use and summarize the general species performance and, according to our results, they appear to be critical for seedling establishment and coexistence. It is remarkable that spatial association of seedlings was not related to differences in SM, which has been considered proxy of dispersal ability (Moles & Westoby, 2004), and that affects also seedling performance and establishment potential (Larson *et al*., 2015). This suggests that performance-related traits, such as SLA or RVR, play a more dominant role in shaping early coexistence patterns than dispersal ability alone. The robustness of these inferences is supported by the large number of species-pair interactions quantified within our plots (>44,300 individuals) and by the consistency of patterns across life stages and spatial scales. In this context, future work combining spatially explicit field approaches with controlled greenhouse experiments manipulating biological soil crust integrity and water availability would represent a valuable next step to isolate the causal mechanisms underlying these assembly rules. Likewise, the two climatically contrasting years analysed here should be interpreted as a test of community assembly responses to sharp environmental contrasts rather than true temporal replication.

Our findings directly contribute to ongoing debates about the predictability of species coexistence from functional traits (Cooksley *et al*., 2024; Westoby, 2025; Levine *et al*., 2025). While previous work suggests that economic functional traits are often weakly correlated with environmental variation (Westoby, 2025), our results provide robust experimental evidence to the contrary. We demonstrate that functional dissimilarity in traits such as SLA, SM, and RVR consistently influences species segregation, especially during wet years and among seedlings, while biological soil crust disturbance modulates trait associations differently at fine and coarse spatial scales. These results highlight that trait-mediated spatial associations are modulated by environmental variation and suggest that spatial organization and coexistence in plant communities reflects, not a fixed response but a predictable dynamic balance between environmental filtering and competitive interactions. Accurately predicting the dynamics of plant community assembly requires considering which traits are implicated, when along the plant life stages they exert influence, and where in space they operate—providing a robust framework for forecasting species coexistence under accelerating environmental change.

## ACKNOWLEDGEMENTS

We thank all participants for their help with feature measurements and digitization of spatial data. EA was supported by a predoctoral fellowship (grant PRE2021-097730). EA, JC and MC were supported by grant PID2020-114851GA-I00 (UNIPER), AE by PID2021-126927NB-I00 (QuerPin), ALL by PID2023-149999NB-I00 (PHYLOFUNKEY), all of them from Agencia Estatal de Investigación (Spanish Government).

## COMPETING INTERESTS

None declared.

## AUTHOR CONTRIBUTIONS

Ezequiel Antorán and Marcelino de la Cruz conceived the ideas and designed methodology; Ana L. Peralta collected the data; Ezequiel Antorán, Marcelino de la Cruz and Joaquin Calatayud analysed the data; Ezequiel Antorán led the writing of the manuscript. All authors contributed substantially to revisions.

## DATA AVAILABILITY

Data is available from the Dryad Digital Repository (https://doi.org/10.5061/dryad.59zw3r2cm). Code will be publicly archived after the acceptance of the paper but would be provided during the review process if required.

## Notes

### Competing Interest Statement

The authors have declared no competing interest.

